# Maize TIR-only Proteins ZmTIR1 and ZmTIR2, but not ZmTIR3 Confer Auto-active Hypersensitive Response Likely by Forming Condensation

**DOI:** 10.1101/2024.12.26.630462

**Authors:** Zi-Xuan Kang, Qi-Dong Ge, Man-Lin Zhang, Chang-Xiao Tang, Man Yang, Shuqing Liu, Yu-Chen Dai, Hua Zhang, Ke Zang, Shengjun Li, Guan-Feng Wang

## Abstract

Nucleotide binding, leucine-rich-repeat (NLR) proteins are the major intracellular receptors for defending against pathogen infection. The recognition between NLRs and pathogen secreted effectors often triggers a localized programmed cell death termed hypersensitive response (HR). Despite significant progresses have been achieved in understanding canonical NLRs with the N-terminal Toll/interleukin-1 receptor (TIR) domains, the molecular mechanisms underlying TIR-only proteins in plant immune responses remain unclear. In this study, we identified six TIR-containing proteins in maize, including three TIR-only proteins. Functional analysis showed that ZmTIR1 and ZmTIR2, but not ZmTIR3, confer autoactive HR when transiently expressed in *N. benthamiana*. The autoactivity conferred by ZmTIR1 and ZmTIR2 depends on EDS1-PAD4-RNL module and their putative NADase activities. Interestingly, ZmTIR1 and ZmTIR2 predominantly localize in the punctate dots and likely form condensation, while ZmTIR3 mainly localizes in the cytoplasm and the nucleus. Two specific amino acids in the BB-loop region were identified to be required for ZmTIR1- and ZmTIR2-mediated condensation formation and auto-HR. Furthermore, *ZmTIR* and *ZmTIR2* are induced by *Cochliobolus heterostrophus*, the causal agent of southern leaf blight (SLB) in maize, and knock-down the expression of *ZmTIR1* or *ZmTIR2* decreased the resistance to SLB in maize. Our study reveals a novel mechanism of monocot TIR-only proteins in maize immune responses.

## INTRODUCTION

Plants utilize a multi-layered surveillance system to defend against pathogen infection. This system includes cell-surface pattern recognition receptors (PRRs) for detecting pathogen-derived conserved molecules to trigger pattern-triggered immunity (PTI) and intracellular immune receptors for recognizing pathogens secreted effectors to confer effector-triggered immunity (ETI). PTI and ETI often act synergistically to recognize pathogens and trigger a series of immune responses, including the accumulation of reactive oxygen species (ROS), a rapid increase of cytosolic calcium (Ca^2+^) and a rapid programmed cell death at the pathogen infection site known as the hypersensitive response (HR) (Sun et al., 2020; Ngou et al., 2021; Yuan et al., 2021). Most intracellular immune receptors encode nucleotide-binding, leucine-rich repeat (NLR) proteins (Kourelis and van der Hoorn, 2018). Canonical NLRs comprise a variable N-terminal domain (NTD), a central NB-ARC (nucleotide-binding adaptors shared by APAF1, certain *R* gene products and CED-4) domain and a C-terminal LRR (leucine-rich repeat) domain (Dangl and Jones, 2001; Sun et al., 2020). The NTDs of plant NLRs mostly encode the Toll/interleukin-1 receptor (TIR), the putative coiled-coil (CC) or the RPW8-type CC (CC_R_) domains, therefore these NLRs are referred as TNLs, CNLs and RNLs, respectively (Shao et al., 2016).

Upon recognition of pathogen effectors, CNLs, for example ZAR1 and Sr35, often form pentameric resistosomes by acting as calcium-permeable channels in triggering plant immunity (Wang et al., 2019; Förderer et al., 2022; Zhao et al., 2022). TNLs often form tetrameric resistosomes, including ROQ1 and RPP1 (Ma et al., 2020; Martin et al., 2020). In TNL resistosomes, the TIR domains exhibit NADase and ADP-ribosylation activities, with NAD^+^ and ATP as substrates, respectively (Horsefield et al., 2019; Wan et al., 2019; Ma et al., 2020; Martin et al., 2020; Jia et al., 2022). TIRs catalyze the substrates to produce small molecules which allosterically activate the heterodimers formed by ENHANCED DISEASE SUSCEPTIBILITY 1 (EDS1)-PHYTOALEXIN DEFICIENT 4 (PAD4) or EDS1-SENESCENCE-ASSOCIATED GENE 101 (SAG101) (Huang et al., 2022; Jia et al., 2022). Once activated, the downstream RNLs, ACTIVATED DISEASE RESISTANCE 1 (ADR1) and N REQUIREMENT GENE 1 (NRG1) interact with EDS1-PAD4 and EDS1-SAG101 heterocomplexes, respectively, and trigger plant immunity (Huang et al., 2022; Jia et al., 2022). Similar to CNLs, RNLs also act by forming pentameric resistosomes which function as Ca^2+^-permeable channels in plant immunity (Jacob et al., 2021; Feehan et al., 2023). EDS1, PAD4 and ADR1 homologs were identified in most seed plant species, while SAG101 and NRG1 homologs were not detected in monocots (Johanndrees et al., 2023). Recently, the Cryo-EM structure of the EDS1-PAD4-ADR1 immune complex was resolved and characterized, and the complex is promoted by small molecules produced by TIR only proteins from arabidopsis, *N. benthamiana*, rice and bacterium (Wang, 2024; Wu, 2024; Yu, 2024).

Except for canonical TNLs, plant genomes also encode TIR-containing proteins that lack canonical TNL domains, including TIR-only proteins, TIR-NB-ARC (TN), and TN fused with tetratricopeptide-like repeats (TPRs) called TNP proteins (Toshchakov and Neuwald, 2020; Johanndrees et al., 2023). Monocot plants do not contain canonical TNLs, but have other TIR-containing proteins (Lapin et al., 2022). TIR-only proteins share conserved downstream signaling components with canonical TNLs in plant immunity. For example, the TIR-only protein Response to HopBA1 (RBA1) in Arabidopsis recognizes effector HopBA1 and trigger EDS1-dependent cell death (Nishimura et al., 2017). Recently, RBA1 and the TIR domain of TNL RPP1 were demonstrated to form phase separation and trigger plant immune responses (Song et al., 2024). Rice TIR-only protein OsTIR plays positive roles in the resistance to *Magnaporthe oryzae* (Wu, 2024). However, the function and molecular mechanisms of TIR-only proteins in plant immune responses, especially in monocots remain less understood.

Here we identified that the maize genome contains three TIR-only proteins, ZmTIR1, ZmTIR2 and ZmTIR3. When transiently expressed in *N. benthamiana*, ZmTIR1 and ZmTIR2, but not ZmTIR3 confer auto-HR which depends on EDS1, PAD4, RNLs and their NADase activities. Interestingly, ZmTIR1 and ZmTIR2 displayed different subcellular localization with ZmTIR3. ZmTIR1 and ZmTIR2 formed condensate-like punctate dots and two key amino acids in the BB-loop region are required for triggering auto-HR and forming condensates. Furthermore, *ZmTIRs* are induced by *Cochliobolus heterostrophus*, the causal agent of southern leaf blight (SLB) in maize. Knock-down the expression of *ZmTIR1* or *ZmTIR2* enhanced the susceptibility to SLB in maize. Our study reveals novel mechanism underlying the action mode of TIR-only proteins in plant immunity.

## RESULTS

### Phylogenetic analysis of plant TIRs

Using hidden Markov models (HMMs) to search the maize genome database (https://www.maizegdb.org), we identified 6 TIR-containing proteins in maize (Table 1), including 3 TIR-only proteins (GRMZM2G319375, GRMZM2G402165, GRMZM2G394261, named as ZmTIR1, ZmTIR2, and ZmTIR3, respectively), 1 TIR-NB protein (GRMZM2G132403, named as ZmTN1), and 2 TNP proteins (GRMZM2G039878 and GRMZM2G067555, named as ZmTNP1 and ZmTNP2, respectively). Phylogenetic analysis showed that ZmTIR1-3 are in the same clade with a few well-known TIRs, such as BdTIR of *Brachypodium distachyum,* OsTIR of rice and Arabidopsis AtRPS4; while ZmTN1 and ZmTNPs are in the same clade with barley HvTIR (Figure 1A).

**Figure 1.**
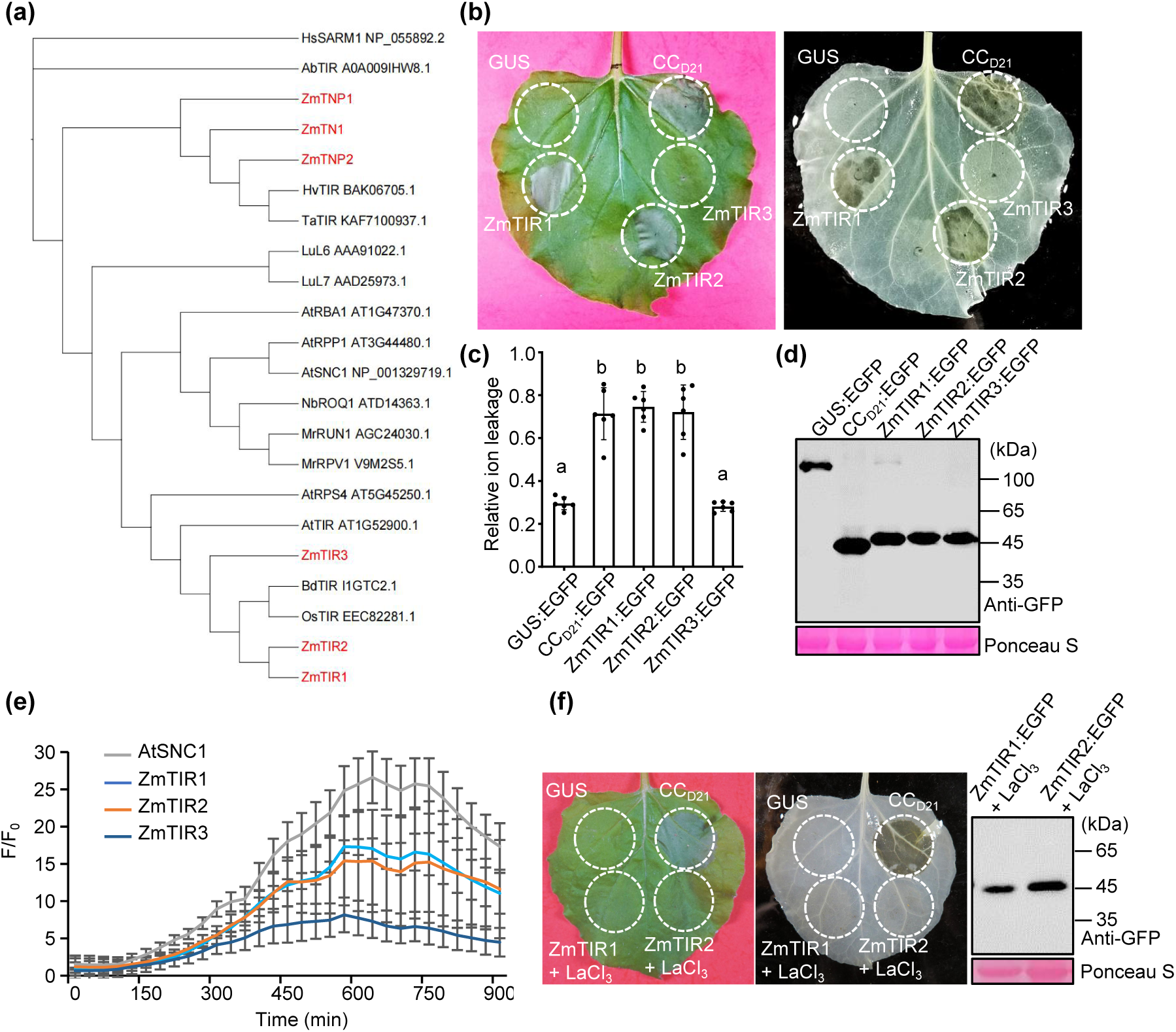
ZmTIR1 and ZmTIR2, but not ZmTIR3 trigger auto-HR in *N. benthamiana*. (a) Phylogenetic analysis of TIR domains from different plant species. Ab: *Acinetobacter baumannii*; At: *Arabidopsis thaliana*; Bd: *Brachypodium distachyon*; Hv: *Hordeum vulgare*; Hs: *Homo sapiens*; Lu: *Linum usitatissimum*; Mr: *Muscadinia rotundifolia*; Nb: *Nicotiana benthamiana*; Os: *Oryza sativa*; Ta: *Triticum aestivum*; Zm: *Zea mays*. (b) GUS, CC_D21_ and ZmTIRs were fused with a C-terminal EGFP and transiently expressed in *N. benthamiana*. A representative leaf (left) was shown at 3 days post-infiltration (dpi), and the same leaf (right) was cleared by ethanol. (c) Ion leakage conductivity was measured at 60 hours post-infiltration (hpi). Different letters (a-b) indicate significant differences (Average ± SE, n > 5, P < 0.05) between samples. (d) Total proteins were extracted from agro-infiltrated *N. benthamiana* leaves at 36 hpi. The expression of different proteins was detected with anti-GFP. Rubisco stained by Ponceau S was used as an equal loading control for protein samples. (e) Time course measurement of ZmTIR-mediated Ca^2+^ influx in GCaMP3-transgenic *N. benthamiana*. (f) The Ca^2+^ inhibitors LaCl_3_ blocked ZmTIR-mediated auto-HR in *N. benthamiana*. 2 mM LaCl_3_ was supplemented in the infiltration buffer before infiltration into *N. benthamiana*, and samples were collected for detecting the protein expression by anti-GFP at 36 hpi. The experiments were repeated at least three times with similar results.

**Table 1.** Information summary of ZmTIRs. The TIR-containing genes in maize.

| Name | Gene ID(v3/v4) | Location | Domain(s) |
| --- | --- | --- | --- |
| ZmTIR1 | GRMZM2G394261/Zm00001d021820 | Chr7:163678563-163680179 | TIR |
| ZmTIR2 | GRMZM2G319375/Zm00001d006617 | Chr2:212688199-212689120 | TIR |
| ZmTIR3 | GRMZM2G402165/Zm00001d007761 | Chr2:239168050-239168955 | TIR |
| ZmTN1 | GRMZM2G132403/Zm00001d053244 | Chr4:221728680-221731688 | TIR-NBARC |
| ZmTNP1 | GRMZM2G039878/Zm00001d052949 | Chr4:205988263-205990716 | TIR-NBARC-TPRs |
| ZmTNP1 | GRMZM2G067555/Zm00001d052845 | Chr4:202430132-202439991 | TIR-NBARC-TPRs |

### ZmTIR1 and ZmTIR2, but not ZmTIR3 confer autoactive HR when transiently expressed in *N. benthamiana*

TIR proteins often confer autoactive HR (auto-HR) when transiently expressed in *N. benthamiana*. To explore whether the maize TIR-only proteins confer auto-HR, we cloned ZmTIR1-3 encoding genes and fused them with the enhanced green fluorescent protein (EGFP) to generate ZmTIRs:EGFP. When transiently expressed in *N. benthamiana* via Agrobacteria, ZmTIR1 and ZmTIR2, but not ZmTIR3 displayed auto-HR at 3 days post-infiltration (dpi) (Figure 1B). Consistent with our previous studies (Wang et al., 2015a; Wang et al., 2015b), the CC domain of the CNL protein Rp1-D21 fused with EGFP (CC_D21_:EGFP) conferred auto-HR and GUS:EGFP did not display auto-HR, which were used as positive and negative controls, respectively (Figure 1B). Consistent with our visual observations, ZmTIR1:EGFP and ZmTIR2:EGFP significantly increased the levels of ion leakage conductivity, which was similar to those caused by CC_D21_:EGFP; while ZmTIR3 exhibited comparable levels of ion leakage conductivity compared to GUS:EGFP (Figure 1C). Immunoblot by anti-GFP antibody confirmed that all proteins had substantial levels of protein accumulation (Figure 1D).

Calcium plays important roles in NLR-mediated HR (Bi et al., 2021; Förderer et al., 2022; Köster et al., 2022). We transiently expressed ZmTIRs in a transgenic *N. benthamiana* line expressing GCaMP3, a single-wavelength fluorescent Ca^2+^ indicator (DeFalco et al., 2017). This system has been widely used for monitor Ca^2+^ signaling in plants in response to biotic stresses or NLR-mediated HR (Jacob et al., 2021; Wang et al., 2021). It was reported that Arabidopsis TNL protein AtSNC1 confers auto-HR and elicits Ca^2+^ levels (Wang et al., 2023), therefore it was used as a positive control. The results showed that ZmTIR1 and ZmTIR2 triggered high levels of Ca^2+^ signaling, though less than AtSNC1, but significantly higher than that caused by ZmTIR3 (Figure 1E). We further treated leaves infiltrated by ZmTIR1 and ZmTIR2 with calcium chelating agent LaCl_3_ and found that LaCl_3_ inhibited ZmTIR1- and ZmTIR2-mediated auto-HR (Figure 1F). These results indicated that calcium is critical for ZmTIR1- and ZmTIR2-mediated auto-HR.

### ZmTIR1- and ZmTIR2-mediated auto-HR depends on EDS1, PAD4 and ADR1/NRG1, but not BAK1 and SOBIR1

EDS1-PAD4-ADR1 and EDS1-SAG101-NRG1 modules are key components acting at the downstream of TIR-mediated immunity (Pruitt et al., 2021). To determine whether ZmTIR1- and ZmTIR2-mediated auto-HR depends on these components, we infiltrated them into the *pad4*, *eds1*, *eds1/pad4/sag101a/sag101b (epss)* mutants and the RNL mutants (*adr1*, *nrg1* and *adr1/nrg1* double mutant) in *N. benthamiana* background (Qi et al., 2018; Lapin et al., 2019; Prautsch et al., 2022; Zonnchen et al., 2022). The results showed that ZmTIR1- and ZmTIR2-mediated HR was abolished in these mutants, while CC_D21_:EGFP still confers strong auto-HR in these mutants (Figure 2A, 2B), indicating that these components are required for ZmTIR1- and ZmTIR2-mediated auto-HR.

**Figure 2.**
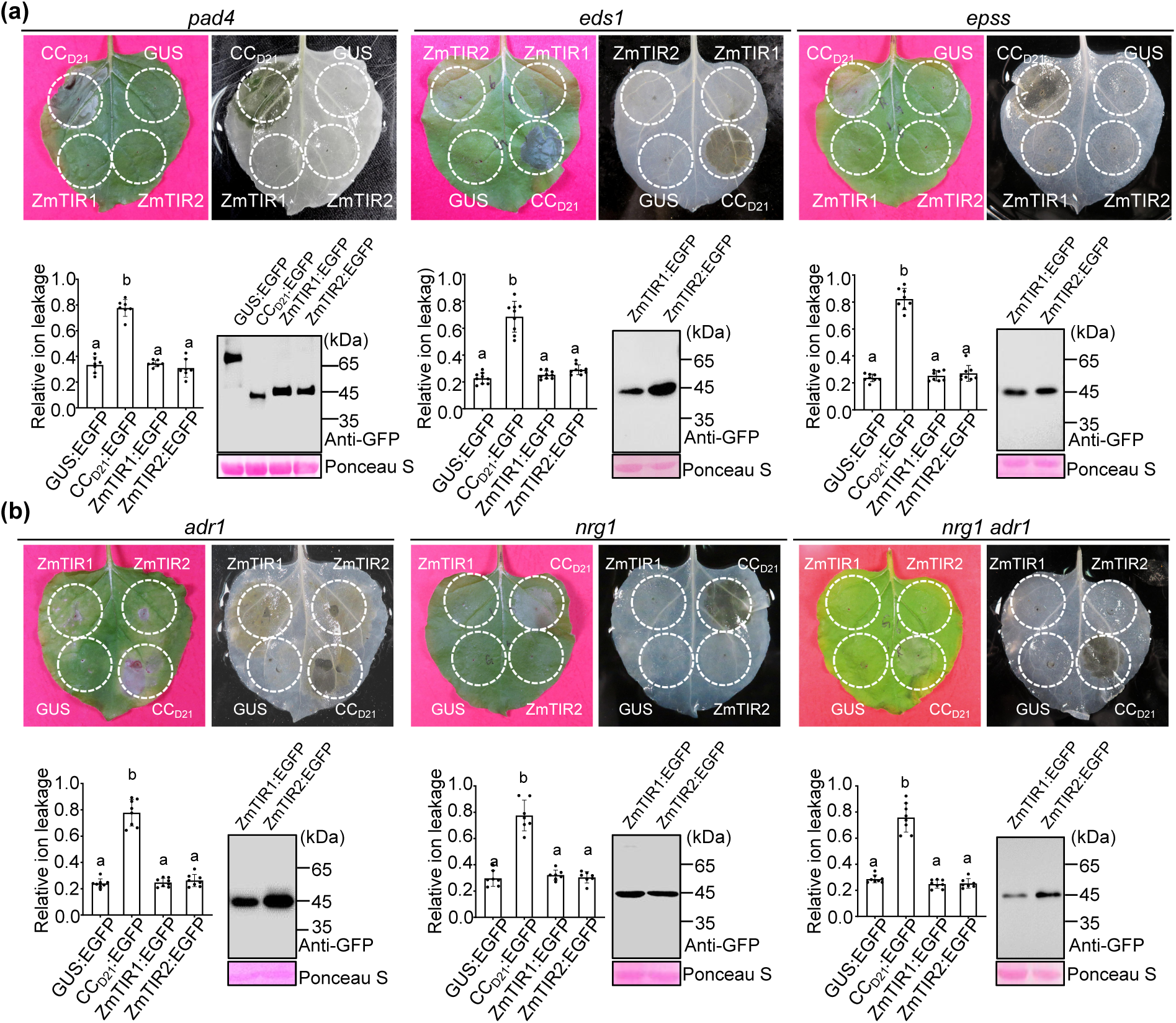
ZmTIR1- and ZmTIR2-mediated auto-HR depends on EDS1-PAD4-RNL module. (a) GUS, CCD21 and ZmTIRs were transiently expressed in *N. benthamiana* mutants *eds1, pad4* and *epss*. A representative leaf (left) was shown at 3 dpi, and the same leaf (right) was cleared by ethanol. (b) GUS, CCD21 and ZmTIRs were transiently expressed in *N. benthamiana* mutants *adr1, nrg1* and *adr1nrg1*. A representative leaf (left) was shown at 3 dpi, and the same leaf (right) was cleared by ethanol. Ion leakage conductivity was measured at 60 hpi. Different letters (a-c) indicate significant differences (Average ± SE, n > 5, P < 0.05) between samples. Total proteins were extracted from agro-infiltrated *N. benthamiana* leaves at 36 hpi. The expression of different proteins was detected with anti-GFP. Rubisco stained by Ponceau S was used as an equal loading control for protein samples.

BAK1 and SOBIR1 are well-known co-receptors for PRRs RLK- and RLP-mediated PTI, respectively (Tian et al., 2021). To determine whether ZmTIR1- and ZmTIR2-mediated auto-HR depends on BAK1 and SOBIR1, we infiltrated them into the *bak1* and *sobir1* mutants in *N. benthamiana* background (Wang et al., 2018; Wang et al., 2022). Not surprisingly, ZmTIR1 and ZmTIR2 still conferred auto-HR in *bak1* and *sobir1* mutants (Figure S1), indicating that BAK1 and SOBIR1 are not required for ZmTIR1- and ZmTIR2-mediated auto-HR.

### The putative NADase activity of ZmTIR1 and ZmTIR2 is required for their auto-activities

TIR proteins exhibit NADase and 2’,3’-cAMP/cGMP synthetase activities and both activities are required for TIR- and TNL-mediated defense responses. The catalytic residues Glu (E) and Cys (C) in αC motif are required for NADase and synthetase activities, respectively, in TIR domains (Horsefield et al., 2019; Wan et al., 2019; Ma et al., 2020; Yu et al., 2022). We found that these residues are conserved in TIRs from different plant species, including ZmTIRs (Figure 3A). Molecular docking demonstrated that ZmTIR1-3 had similar structure to the TIR domains from ROQ1, RPP1 and RBA1 (Figure 3B). To explore the function of the putative enzymatic activities in ZmTIR1- and ZmTIR2-mediated HR, we generated the corresponding mutations in ZmTIR1 and ZmTIR2. When transiently expressed in *N. benthamiana*, ZmTIR1(E134A) and ZmTIR2(E139A) which presumably disrupt the NADase activity abolished auto-HR, while ZmTIR1(C131A) and ZmTIR2(C136A) which presumably disrupt the synthetase activity had no obvious effects (Figure 3C-3E). Furthermore, in the GCaMP3 transgenic *N. benthamiana* line, the Ca^2+^ signaling in ZmTIR1(E134A) and ZmTIR2(E139A) was significantly reduced than that in ZmTIR1 and ZmTIR2 (Figure 3G). These results indicated that the putative NADase activity is required for ZmTIR1- and ZmTIR2-mediated auto-HR.

**Figure 3.**
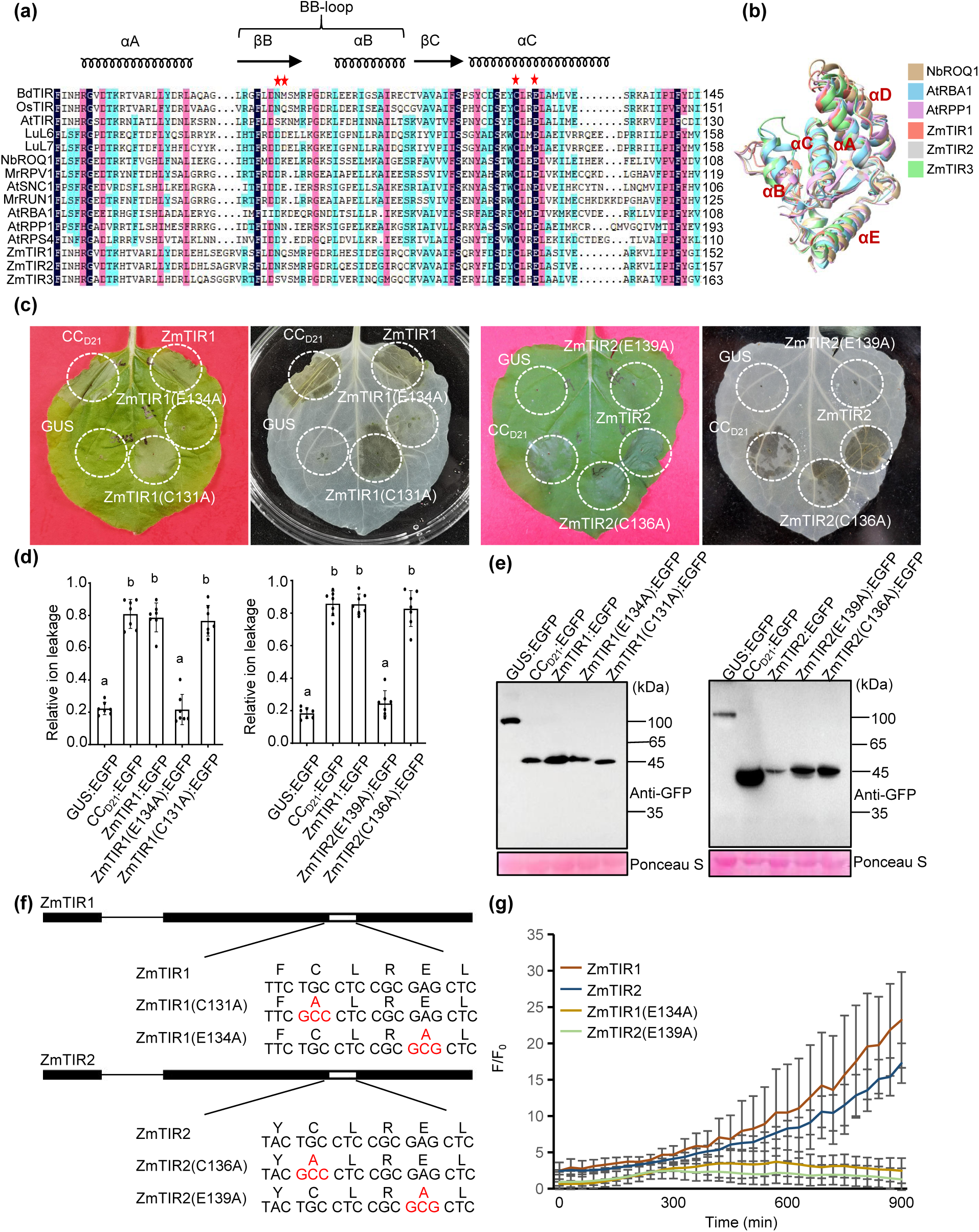
The NADase activity is required for ZmTIR1- and ZmTIR2-mediated auto-HR. (a) Multiple sequence alignment of TIR proteins from different plant species. The conserved Cys (C) and Glu (E) residues at position 131 and 134 of ZmTIR1 are labeled with red stars. (b) The predicted structure of TIR proteins from maize, *N. benthamiana* and Arabidopsis. (c) ZmTIR1, ZmTIR2 and their mutant derivatives were transiently expressed in *N. benthamiana*. The representative leaves were shown at 3 dpi. (d) Ion leakage conductivity was measured at 60 hpi. Different letters (a-b) indicate significant differences (Average ± SE, n > 5, P < 0.05) between samples. (e) Total proteins were extracted from agro-infiltrated *N. benthamiana1* leaves at 36 hpi. The expression of different proteins was detected with anti-GFP. Rubisco stained by Ponceau S was used as an equal loading control for protein samples. (f) The diagram of mutant derivatives of *ZmTIR1* and *ZmTIR2*. (g) Time course measurement of ZmTIR- and the derivative-mediated Ca2+ influx in GCaMP3-transgenic *N. benthamiana*.

### The BB-loop is required for the autoactivity mediated by ZmTIR1 and ZmTIR2

BB-loop is important for the function of some TIRs, and the mutations in the BB-loop region abolish the auto-HR conferred by BdTIR1, RBA1 and RPS4 (Bayless et al., 2023). However, the BB-loop mutation in the prokaryotic TIR-containing protein ThsB enhanced the autoactivity (Bayless et al., 2023). To investigate the role of the BB-loop in ZmTIR-mediated HR, we generated the BB-loop mutations in ZmTIR1 and ZmTIR2 to produce ZmTIR1(bl) and ZmTIR2(bl) (Figure 4A). When infiltrated into *N. benthamiana*, ZmTIR1(bl) and ZmTIR2(bl) abolished the auto-HR (Figure 4B). We further mutated two amino acids (NQ in ZmTIR1, NK in ZmTIR2) in the BB-loop region to generate ZmTIR1(NQ/GG) and ZmTIR2(NK/GG), and found that both mutants abolished the auto-HR, which was similar to the effects caused by ZmTIR1(bl) and ZmTIR2(bl) (Figure 4A, 4B). These results indicated that the BB-loop region, especially the NQ/NK residues in ZmTIR1/ZmTIR2 are required for their autoactivity.

**Figure 4.**
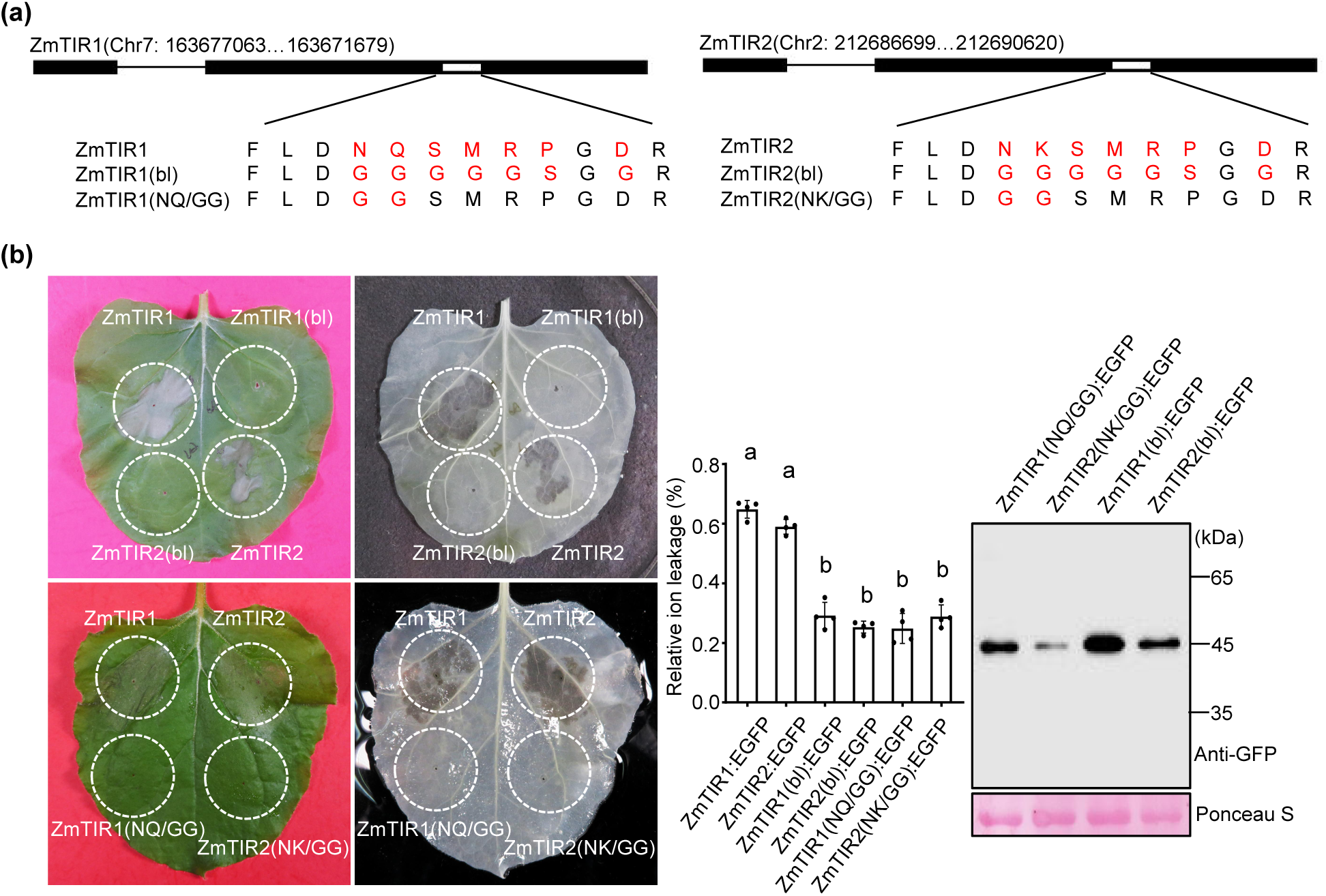
The BB-loop is required for ZmTIR1- and ZmTIR2-mediated auto-HR. (a) The diagram of BB-loop mutant derivatives of *ZmTIR1* and *ZmTIR2*. (b) ZmTIR1, ZmTIR2 and their BB-loop mutant derivatives were transiently expressed in *N. benthamiana*. The representative leaves were shown at 3 dpi. Ion leakage conductivity was measured at 60 hpi. Different letters (a-b) indicate significant differences (Average ± SE, n > 5, P < 0.05) between samples. Total proteins were extracted from agro-infiltrated *N. benthamiana* leaves at 36 hpi. The expression of different proteins was detected with anti-GFP.

### ZmTIR1, ZmTIR2 and ZmTIR3 self-associate

TIRs often self-associate to form oligomers for function. We observed that ZmTIR1, ZmTIR2 and ZmTIR3 can self-associate as detected by split LUC and yeast two-hybrid (Figure 5A, 5B). Interestingly, when detected by bimolecular fluorescence complementation (BiFC) assays, ZmTIR1 and ZmTIR2 self-associated in the punctate dots while ZmTIR3 formed homomers in the cytoplasm and the nucleus (Figure 5C). Intriguingly, the NADase mutants ZmTIR1(E134A) and ZmTIR2(E139A) and the BB-loop mutant ZmTIR2(NK/GG) still self-associated when detected by co-immunoprecipitation (Co-IP) assays (Figure S2).

**Figure 5.**
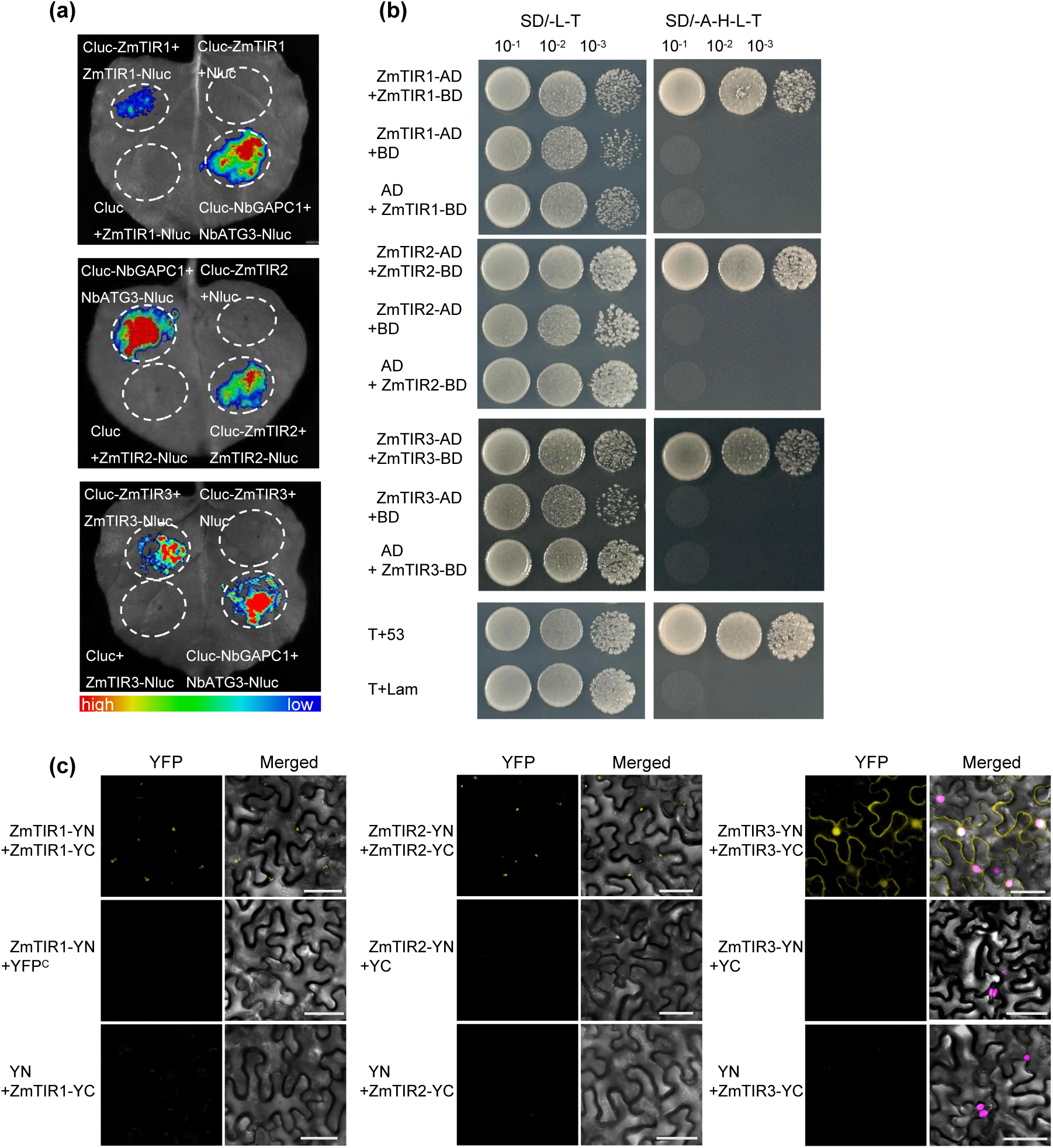
ZmTIRs self-associate. (a) Self-association of ZmTIRs were detected by split luciferase assays in *N. benthamiana*. A representative image was displayed at 60 hrs. (b) Yeast two hybridization (Y2H) assay was used to investigate the self-association of ZmTIRs. T + 53 and T + Lam are used as positive and negative controls, respectively. (c) Bimolecular fluorescence complementation (BiFC) assays were used to investigate the self-association of ZmTIRs. ZmTIRs were constructed into split YFP-derived YN and YC vectors, and they were co-expressed into *N. benthamiana*, and the fluorescence was observed at 40 hpi by confocal microscopy. The scale bar represents 50 µm.

### ZmTIR1 and ZmTIR2 have different subcellular localization compared to ZmTIR3

To investigate the subcellular localization of ZmTIRs, we transiently expressed ZmTIRs:EGFP in a transgenic *N. benthamiana* line carrying the nuclear marker H2B:TaqRFP for confocal observation (Martin et al., 2009). ZmTIR1 and ZmTIR2 mainly localized in the punctate dots, with some weak signals in the nucleus and cytoplasm, while ZmTIR3 mainly localized in the nucleus and cytoplasm (Figure 6A). We further verify the results in maize protoplasts by co-transformation of GFP-tagged ZmTIRs (ZmTIRs:GFP) and the nuclear marker H2A:mcherry. Consistent with the results in *N. benthamiana*, ZmTIR1 and ZmTIR2 mainly displayed a punctate pattern of localization in protoplasts, with weak signals in the nucleus and cytoplasm; while ZmTIR3 was evenly distributed in the cells (Figure 6B). Moreover, we performed biochemical protein fractionation assays to verify the results. For the microsomal fractionation assays, ZmTIR1, ZmTIR2 and ZmTIR3 mainly localized in the cytoplasm, with minor in the membrane (Figure 6C). For the nuclear fractionation assays, ZmTIR1, ZmTIR2 and ZmTIR3 all have nuclear enrichment, except for the soluble fraction (Figure 6D). These results indicated that ZmTIR1 and ZmTIR2 mainly displayed a punctate distribution pattern in the cytoplasm, with minor in the nucleus and the membrane; ZmTIR3 predominantly localized in the cytoplasm and the nucleus, with minor in the membrane.

**Figure 6.**
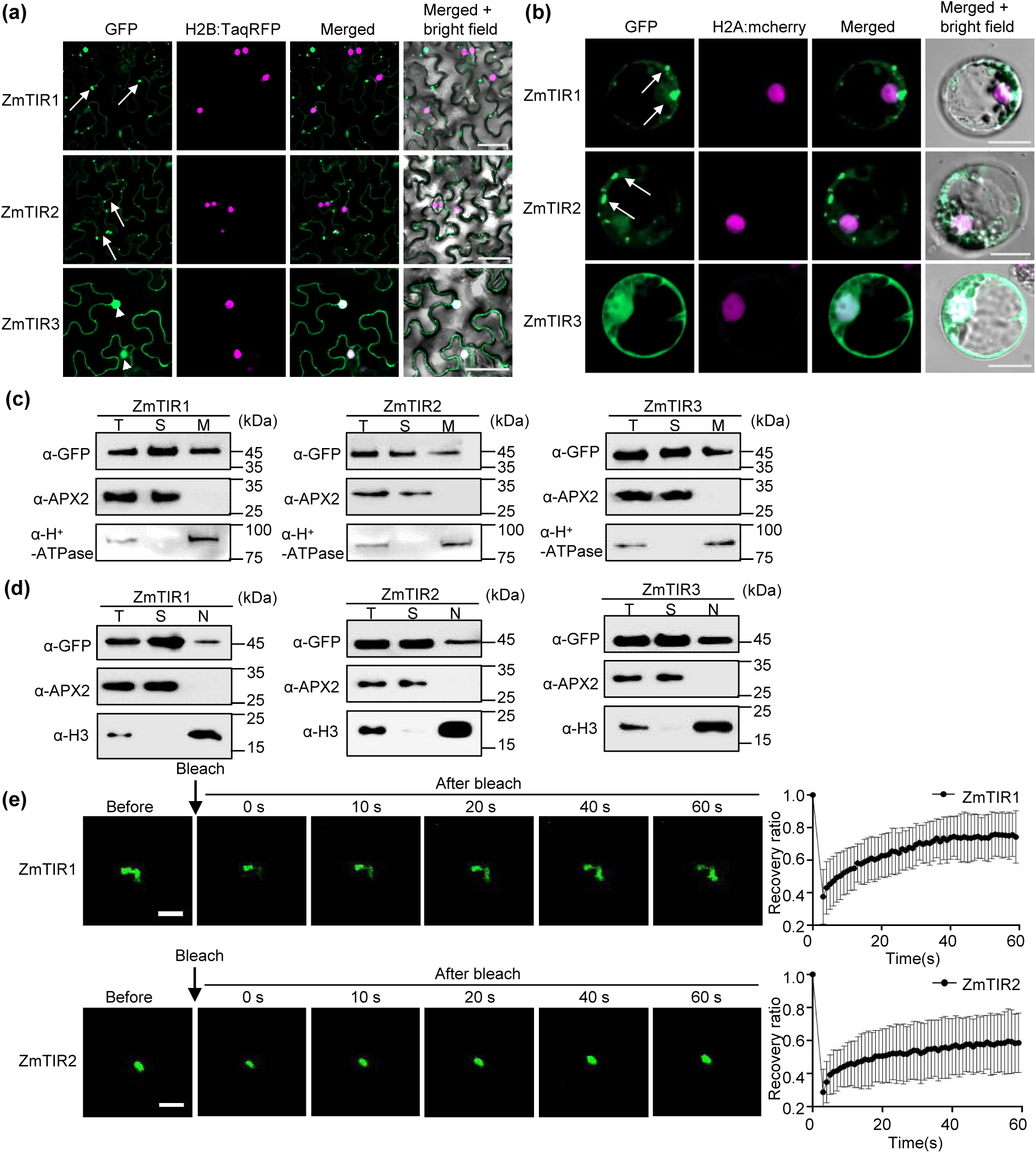
ZmTIR1 and ZmTIR2 exhibit different subcellular localization patterns with ZmTIR3 in *N. benthamiana* and maize protoplasts. (a) ZmTIRs:EGFP were infiltrated into *N. benthamiana* harboring the nuclear marker H2B:TaqRFP. Confocal images were taken at 36 hpi. The positions of the nucleus and the punctate dots were labeled by arrowheads and arrows, respectively. The scale bar represents 50 µm. (b) ZmTIRs:GFP were transiently co-expressed with the nuclear marker H2A:mcherry in maize protoplast. Confocal images were taken at 18 hpi. Arrows indicate the positions of the punctate dots. The scale bar represents 20 µm. (c) Protein microsomal fractionation assays. ZmTIRs:EGFP were transiently expressed in *N. benthamiana* and total protein (T), soluble (S) and microsomal (M) fractions were extracted at 48 hpi. Anti-APX2 and anti-H^+^-ATPase were used as soluble and microsomal markers, respectively. (d) Protein nuclear fractionation assays. ZmTIRs:EGFP were transiently expressed in *N. benthamiana* and total protein (T), soluble (S) and nuclear (N) fractions were extracted at 48 hpi. Anti-APX2 and anti-H3 were used as soluble and nuclear markers, respectively. The experiments were repeated three times with similar results. (e) ZmTIR1 and ZmTIR2 form condensates. ZmTIR1:EGFP and ZmTIR2:EGFP were transiently expressed in *N. benthamiana*, and the confocal images for the dynamics analysis of FRAP were taken at 40 hpi. The scale bar represents 20 µm. The recovery curves of FRAP assays were shown on the right. Data are presented as mean ± SD (n >7 independent biological samples).

### ZmTIR1 and ZmTIR2 form condensates

The punctate localization in the cytoplasm of ZmTIR1 and ZmTIR2 is reminiscent of the formation of condensates or phase separation (Zavaliev et al., 2020; Song et al., 2024). We therefore use PONDR to predict the intrinsically disordered regions (IDRs) in ZmTIRs and found that all the three ZmTIRs displayed IDRs in the N-termini, however, ZmTIR1 and ZmTIR2, but not ZmTIR3 displayed IDRs in the BB-loop regions (Figure S3A). To verify the hypothesis, we transiently expressed ZmTIR1:EGFP and ZmTIR2:EGFP in *N. benthamiana* and performed fluorescence recovery after photobleaching (FRAP) assays. The results showed that both ZmTIR1:EGFP and ZmTIR2:EGFP displayed fluorescence recovery (Figure 6E), suggesting that ZmTIR1 and ZmTIR2 form phase separation.

Intriguingly, the BB-loop mutants ZmTIR1(NQ/GG) and ZmTIR2(NK/GG) predominantly localized in the cytoplasm and nucleus, without punctate dots (Figure S3B). However, the catalytic dead mutants ZmTIR1(E134A) and ZmTIR2(E139A) still formed condensate-like puncta (Figure S3B).

### ZmTIR1 and ZmTIR2 play critical roles in the resistance to maize SLB

To explore the biological relevance of ZmTIRs in maize disease resistance, we investigated the transcript levels of *ZmTIRs* in maize seedling inoculated with *Cochliobolus heterostrophus* (*C.h*), the causal agent of SLB. We found that *ZmTIR1* and *ZmTIR2* which conferred auto-HR were significantly induced by *C.h* at 24 hours post-inoculation (hpi) and dropped at 48 hpi (Figure 7A), suggesting that they might act in maize disease resistance to SLB. To determine the function of ZmTIRs in maize resistance to SLB, we used cucumber mosaic virus (CMV)-mediated virus-induced gene silencing (VIGS) to knock down the expression of *ZmTIRs*. *ZmTIR1* and *ZmTIR2* had more than 85% sequence similarity at the nucleotide level, so we designed a conserved fragment between *ZmTIR1* and *ZmTIR2* to generate the construct *CMV:ZmTIR1/2* for silencing *ZmTIR1* and *ZmTIR2* simultaneously. The transcript levels of *ZmTIR1* and *ZmTIR2* were very low in the normal condition (maizeGDB and Figure 7A), therefore we detected the silencing effect of *ZmTIR1s* by qRT-PCR for samples challenged with *C. heterostrophus*. Compared to negative control *CMV:N81*, the transcript levels of *ZmTIR1 and ZmTIR2* were reduced in *CMV:ZmTIR1/2* while the transcript level of *ZmTIR3* had no obvious effect (Figure 7B), indicating that *ZmTIR1 and ZmTIR2* were silenced. After inoculation with *C.h*, the disease symptoms were significantly larger on leaves infected by *CMV:ZmTIR1/2* than those by the negative control (Figure 7C-D). These results suggested that ZmTIR1 and ZmTIR2 positively regulate the resistance to SLB in maize.

**Figure 7.**
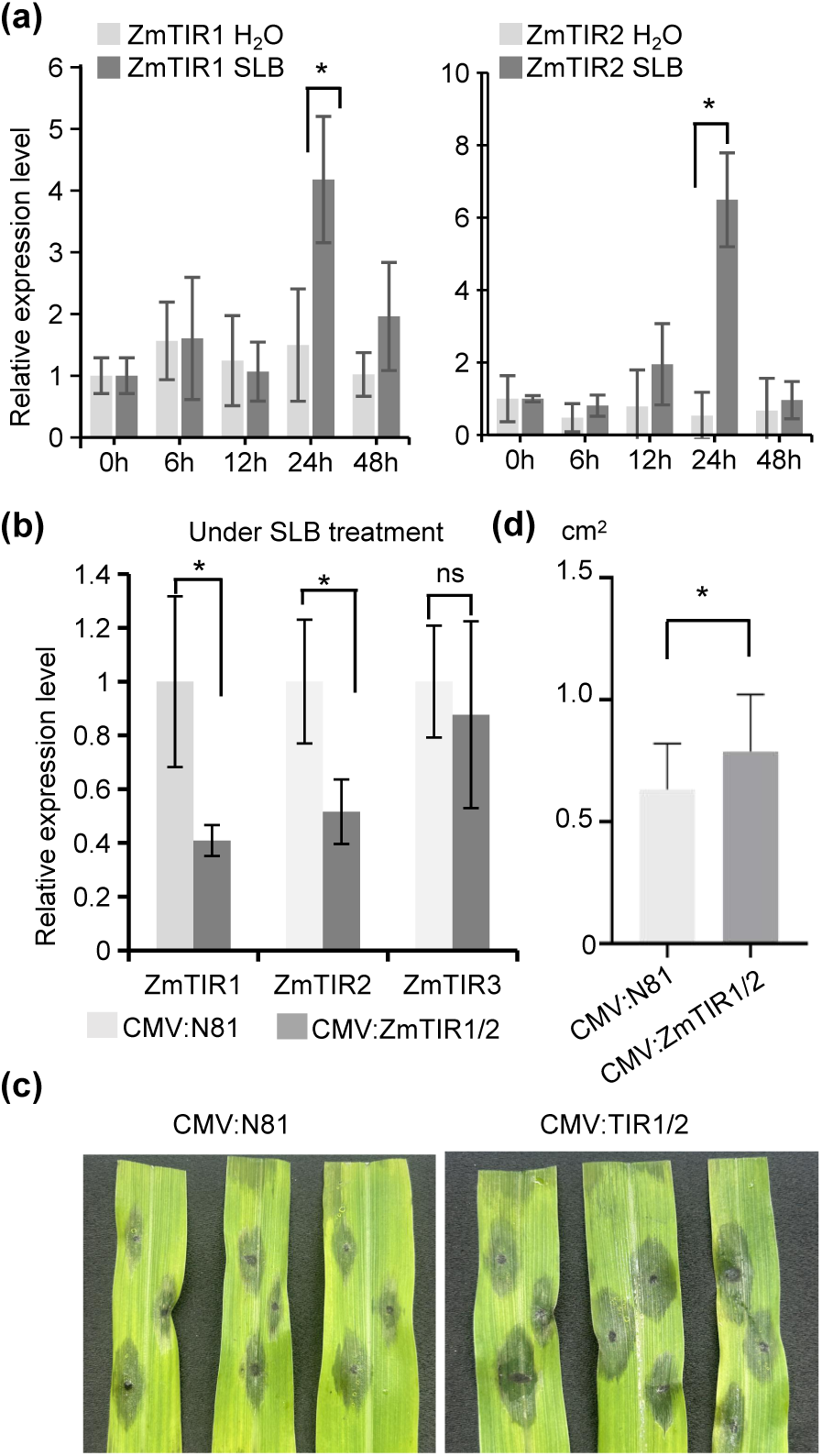
ZmTIRs play critical roles in the resistance to southern leaf blight. (a) The transcript levels of *ZmTIRs* were detected by qRT-PCR in maize inbred line inoculated by *Cochliobolus heterostrophus* at the indicated time points. *ZmActin* was used as an internal control. Data are shown as means ± SE (n = 6). Significant difference between samples (\*\*\**p* < 0.001; \*\**p* < 0.01; ns, not significant) was determined by Student’s *t*-test. (b) The transcript levels of *ZmTIRs* were detected by qRT-PCR when *ZmTIRs* were silenced by CMV-VIGS. (c) The relative disease area of SLB when *ZmTIRs* were silenced by CMV-VIGS. Data are shown as means ± SE (n > 30). Significant difference between samples (\**p* < 0.05; ns: no significant difference) was determined by Student’s *t*-test. (d) The disease phenotypes of SLB when *ZmTIRs* were silenced by CMV-VIGS. Images were taken at 3 dpi. The experiments were repeated three times with similar results.

## DISCUSSION

The immune responses and cell death mediated by TIR domains are largely conserved in plants (Lapin et al., 2022). Curiously, monocots do not encode canonical TNLs, but TIR-only proteins that can cross-activate TIR-immune pathways in dicot. However, the immune function and molecular mechanisms of TIR-only proteins in monocots remain less characterized. In monocot *Brachypodium distachyon*, the TIR-only protein BdTIR triggers EDS1-dependent auto-HR in *N. benthamiana*, but its immune function in *B. distachyon* has not been demonstrated (Wan et al., 2019). Though RBA1 and RPP1-TIR are in the different clades with BdTIR based on the phylogenetic analyses (Figure 1A), all of them confer EDS1-dependent cell death responses (Wan et al., 2019; Johanndrees et al., 2023; Song et al., 2024). Recently, a maize TIR-containing protein TNP has been reported to elicit EDS1-independent cell death in *Nicotiana tabacum* (Johanndrees et al., 2023). However, the function of TIR-only proteins in maize remains unknown. Interestingly, we showed that ZmTIR1, ZmTIR2 and ZmTIR3 are classified into the same clade with BdTIR (Figure 1A), however, ZmTIR1 and ZmTIR2, but not ZmTIR3 confer auto-HR, suggesting they have functional divergence in plant defense responses. RBA1 and RPP1-TIR form phase separation and trigger plant immune responses in Arabidopsis (Song et al., 2024). Similar to RBA1 and RPP1-TIR, ZmTIR1 and ZmTIR2, but not ZmTIR3 formed condensates (Figure 6E). The formation of condensates or the cell death activity of TIR proteins strongly depend on their concentrations (Nishimura et al., 2017; Shin and Brangwynne, 2017; Song et al., 2024). Since ZmTIR1, ZmTIR2 and ZmTIR3 are transiently expressed with the same strong promoter 35S, and they were expressed at largely comparable levels (Figure 1D), the different HR phenotype conferred by them is not caused by the difference of the protein concentrations. Several TIR domain-mediated auto-HR requires EDS1, SAG101b and NRG1, but not PAD4, SAG101a and ADR1 (Qi et al., 2018; Gantner et al., 2019; Lapin et al., 2019; Chen et al., 2024). Here we showed that ZmTIR1 and ZmTIR2 triggered auto-HR not only depends on EDS1 and NRG1, but also PAD4 and ADR1, indicating that these components act at downstream of ZmTIR1 and ZmTIR2.

The NADase activity is required for TIR domain protein-mediated cell death (Wan et al., 2019; Ma et al., 2020). Not surprisingly, when the conserved residues required for NADase activity were mutated, ZmTIR1- and ZmTIR2-mediated auto-HR were also abolished (Figure 3). However, these mutants did not affect the punctate localization and the self-association of ZmTIR1 and ZmTIR2 (Figures S2, S3B), suggesting that the NADase activity is not required for the formation of condensates. These results are consistent with the corresponding mutation in the TIR protein RBA1 (Song et al., 2024), indicating the functional conservation in different TIR proteins. Except for NADase activity, TIR proteins also exhibit 2’, 3’-cAMP/cGMP synthetase activity by hydrolyzing RNA/DNA and the conserved catalytic residue Cys (C) in TNLs L7 and RBA1 is required for this activity (Yu et al., 2022). Mutations of Cys (C) in TIR-only RBA1, TNLs L6 and RPV1 suppressed the auto-HR in *N. benthamiana* (Bernoux et al., 2011; Williams et al., 2016; Yu et al., 2022). However, when the corresponding Cys (C) were mutated in ZmTIR1 and ZmTIR2, they had no obvious effects on auto-HR (Figure 3), indicating that the putative synthetase activity is not required for ZmTIR1- and ZmTIR2-mediated HR. Similar to our results, the equivalent mutations affecting the putative synthetase activity in the full-length TNL SNC1 do not suppress SNC1-mediated auto-HR both in *N. benthamiana* and Arabidopsis (Tian et al., 2022). The results from full-length TNL SNC1 together with those from our TIR-only ZmTIR1 and ZmTIR2 raise the cautions about the biological importance of synthetase activity in plant immune responses.

BB-loop plays important roles in TIR-mediated immune responses (Ma et al., 2020; Martin et al., 2020). Since ZmTIR1 and ZmTIR2 displayed different HR phenotypes with ZmTIR3, we compared their amino acid sequences and identified two novel key amino acids (NQ in ZmTIR1 and NK in ZmTIR2) in the BB-loop regions. ZmTIR1(NQ/GG) and ZmTIR2(NK/GG) mutants nearly abolished the auto-HR (Figure 4) and greatly reduced the localization in the punctate dots (Figure S3), supporting an essential role of the punctate structures facilitated by the BB-loop in ZmTIR1/2-mediated auto-HR. BB-loop plays important roles in driving phase separation and the NAD^+^-hydrolysing activity of TIR proteins RPP1-TIR, RBA1 and TX14 (Song et al., 2024). We speculated that the BB-loop in ZmTIR1 and ZmTIR2 might have conserved roles in phase separation and NADase activities with other TIR proteins, which needs to be further investigated in the future. RPP1-TIR exhibits nuclear and peri-nuclear puncta, while TX14 exhibits nucleocytoplasmic puncta (Song et al., 2024). RBA1 and TX21 display as cytoplasmic puncta (Nandety et al., 2013; Nishimura et al., 2017; Song et al., 2024). We found that ZmTIR1 and ZmTIR2 exhibit a punctate pattern in the cytoplasm, with minor localization in the membrane and nucleus, while ZmTIR3 does not form puncta. These results indicated that TIR proteins have diverse subcellular localizations, which might differentiate their different functions in defense responses. Oligomer formation is required for TIR activities. We found that all the three ZmTIRs can self-associate, indicating that the inability for ZmTIR3 to induce HR is not related to self-association.

Bimolecular condensates or phase separation play dual roles in plant immunity. Phase separation formed by TIR proteins promotes TIR NADase activity and triggers immune responses, e.g. auto-HR (Song et al., 2024), while NPR1 condensates promote cell survival during plant immunity (Zavaliev et al., 2020). These results suggest phase separation plays a broad role in plant immune responses. In this study, we found that ZmTIR1 and ZmTIR2, but not ZmTIR3 form condensates and confers auto-HR when they were transiently expressed in *N. benthamiana* under strong promoter 35S. We speculated that the condensates formed by ZmTIR1 or ZmTIR2 might affect the transcriptional reprogramming of genes in defense responses, therefore ZmTIR1/ZmTIR2 exhibited different HR phenotype with ZmTIR3. Interestingly, the genes encoding ZmTIR1 and ZmTIR2 are barely detected in normal conditions without any treatments, however they were significantly induced by *C.h* (Figure 7A). When *ZmTIR1* and *ZmTIR2* were knocked down by VIGS, the plants were more susceptible to SLB, indicating that ZmTIR1 and ZmTIR2 play important roles in the resistance to SLB. These results also indicated that ZmTIR-mediated auto-HR and disease resistance can be un-coupled. We have tried to verify the results by getting the transgenic overexpression lines but it failed. It might be caused by the autoimmunity when ZmTIR1 and ZmTIR2 were driven by constitutive strong promoter ubiquintin. In the future, the phenotypes in the lines with ZmTIR1 and ZmTIR2 driven by their native promoters or in the genome-editing lines generated by CRISPR/Cas9 should be further investigated. This study laid the basis for further exploring the function and molecular mechanisms of ZmTIRs in maize immunity.

## EXPERIMENTAL PROCEDURES

### Plant Materials and Growth Conditions

Wild type *N. benthamiana*, *eds1*, *pad4*, *epss*, *adr1*, *nrg1*, *nrg1adr1*, *sorbir1*, *bak1* mutants and the transgenic lines containing H2B:TaqRFP or GCaMP3 were grown in growth chamber at 24°C with 16 hrs light/8 hrs dark. The seedlings of maize inbred line B73 were grown at constant 24 °C with 12 hrs light/12 hrs dark and used for isolating the cDNA of *ZmTIR* genes.

### Plasmid Construction

GUS:EGFP and CC_D21_:EGFP were generated previously (Wang et al., 2015). *ZmTIR* genes were amplified from B73 cDNA with primers listed in Table S1. The PCR products were cloned into pENTR directional TOPO cloning vector (Invitrogen) and then transferred into the destination vector pSITEII-N1-EGFP (Martin et al., 2009) for transient expression in *N. benthamiana* or PUC19-derived vector containing GFP (Gao et al., 2014) for transient assay in maize protoplasts.

### Sequence Alignment and Phylogenetic Analysis

Amino acid sequences from different plant TIRs were aligned using ClustalW (www.ebi.ac.uk). The phylogenetic analysis was performed with MEGA 6.0 software (Tamura et al., 2013). The specific algorithm used for constructing the phylogenetic tree was neighbour-joining with 1000 bootstrap replicates.

### *Agrobacteria*-Mediated Transient Expression in *N. benthamiana*

Different binary vectors were transformed into *Agrobacterium tumefaciens* strain GV3101 (containing pMP90). The details for Agro-bacteria mediated transient expression in *N. benthamiana* were conducted according to our previous studies (Wang and Balint-Kurti, 2015; Wang et al., 2015). Unless otherwise stated, all experiments were performed at least three times with similar results.

### Protein Analysis and Microsomal Fractionation Assays

The details for protein extraction and western blot assays were performed as described in our previous studies (Wang et al., 2015; Wang and Balint-Kurti, 2016; Luan et al., 2021; Sun et al., 2023). Microsomal fractionation assays were conducted based on our previous studies (Wang and Balint-Kurti, 2015; Sun et al., 2023).

### Nuclear Fractionation Assays

Nuclear fractionation assays were conducted based on previous study (Zhao et al., 2010), with minor modification. About 1.0 g leaf tissues were ground in pre-chilled mortars with liquid nitrogen, then add 3 mL of pre-cooling buffer1(1 mM Tris-HCl [pH=7.5], 2 mM Sucrose, 1 mM MgCl_2_, 2 mM KCl, 50% Glycerin, 0.25% Triton X-100, 0.2% MG132, 0.5% β-Mercaptoethanol, 1% MG132). The resulting lysate was filtered with double layers of Miracloth and 100µl supernatant was retained as T1 (Total protein). The remaining supernatant was centrifuged at 2,500 g for 20 min and the supernatant was gently collected as S1 (the soluble fraction). The precipitate was resuspended in 1 mL of buffer 2 (1 mM Tris-HCl [pH=7.5], 2 mM KCl, 0.2% EDTA, 50% Glycerin, 0.25% Triton X-100, 0.2% MG132) and then left on ice for 10 minutes. The supernatant was centrifuged at 12000 g for 10 min, and the precipitate was resuspended in 50 μl of buffer 1 and as N1 (the nuclear fraction). The above samples were mixed with 2× Laemmli buffer for SDS-PAGE.

### Semi-quantification of Calcium Levels in *N. benthamiana*

ZmTIRs and their derives were transiently expressed in a transgenic *N. benthamiana* line expressing a fluorescent Ca^2+^ indicator GCaMP3, and Ca^2+^ levels were measured by a semi-quantitative method as described previously (DeFalco et al., 2017; Jacob et al., 2021) with minor modification. Briefly, 8 leaf discs (0.8 cm diameter) from different single plants were collected at 30 hai and equilibrated in 100 µl ddH_2_O for 6 h in a 96-well flat bottom black plates (Thermo-Fisher). GCaMP3 fluorescence was recorded for each leaf disc at 30 s intervals by a SpectraMax i3x plate-reader, with excitation at 475 nm (20 nm bandwidth) and emission detection at 525 nm (20 nm bandwidth). Absolute fluorescence values were normalized to the untreated control value as F/F_0_ (F was the fluorescence measured at a given time point and F_0_ was the averaged measurement for water-equilibrated control samples measured at the final resting time point).

### Yeast Two-Hybrid (Y2H) Assay

ZmTIRs were constructed into pGADT7 (AD) and pGBKT7 (BD) vectors (Clontech). Different AD- and BD-derived vectors were co-transformed into yeast strain Y2HGold. The detailed procedures for Y2H assays were conducted based on the manufacturer’s protocol (Clontech).

### Split Luciferase Assays

ZmTIRs were constructed into the N- or C-termini of luciferase (Nluc or Cluc) vectors. Different vectors were transiently co-infiltrated into *N. benthamiana* and samples were collected at 48 hai. After treated with 0.5 mM luciferin, the luminescence signals were recorded using CLINX luminescent imaging system (ChemiScope6000, Clinx Science Instruments).

### BiFC Assays in *N. benthamiana*

ZmTIRs were constructed into split YFP-derived YN and YC vectors pEarleyGate 201 and pEarleyGate 202. Different vectors were transiently co-expressed into *N. benthamiana*, and images were taken at 48 hai by confocal microscopy (LSM 880, Carl Zeiss).

### Transient Expression in Maize Protoplasts

The detailed protocol for isolation maize protoplasts and the transfection methods were conducted as reported previously (Jang and Sheen, 1994; Luan et al., 2021; Sun et al., 2023). The protoplasts were observed by a confocal microscope (LSM 880, Carl Zeiss) at 18 hrs after transfection.

### Confocal Microscopy Observation

Microscopic observation was conducted by a confocal microscope (LSM 880, Carl Zeiss) as described in our previous studies (Wang and Balint-Kurti, 2015; Luan et al., 2021). Confocal images were taken at 40 hai in *N. benthamiana* and 24 hai in maize protoplasts.

### FRAP assays

FRAP of ZmTIRs:GFP condensates in *N. benthamiana* leaf epidermal cells was conducted on an Olympus FluoView™ FV1000 laser scanning confocal microscope with an X60/1.42 oil immersion objective. The punctate dots were photobleached using a 100% laser intensity at the 488 nm wavelength with 12 iterations. The recovery time was recorded every 1 s for 60 s after bleaching. Images were acquired and analyzed using the FV10-ASW4.2 software.

### RNA extraction and qRT-PCR analyses

Total RNA was extracted from leaves of maize seedlings using RNAiso reagent (Takara, http://www.takara.com.cn). First strand cDNA was synthesized using HiScript III RT SuperMix for qRCR (Vazyme, http://www.vazyme.com) according to the manufacturer’s instructions. qRT-PCR analysis was performed with a qTower^3^G real-time PCR system (Analytik Jena) using SYBR qPCR Mix (Vazyme), with *ZmActin* as the reference gene (Sun et al., 2023). The data were calculated using the 2*^-^*^△△^*^CT^* method (Livak and Schmittgen, 2001). The primers are listed in Supplemental Table 1.

### CMV-mediated VIGS in Maize

To silence *ZmTIR* genes in maize, a conserved 175-bp DNA fragment between *ZmTIR1* and *ZmTIR2* was cloned from *ZmTIR1* and constructed into pCMVZ22bN81 to generate pCMV::ZmTIR1/2, and a 155-bp DNA fragment was cloned from *ZmTIR3* and constructed into pCMVZ22bN81 to generate pCMV::ZmTIR3. The primers used are listed in Supplemental Table 1. The details for CMV-VIGS were performed according to previous studies (Wang et al., 2016; Li et al., 2021) with minor modification. Different CMV constructs were transformed into *A. tumefaciens* strain GV3101. Agrobacteria harboring CMV RNAs 1, 2, and 3 constructs or their derivatives were equally mixed and agro-infilatrated into *N. benthamiana*. Samples were collected at 4 dpi and grounded in inoculation buffer (20 mM Na2HPO4-NaH2PO4 buffer, pH 7.2) with a 1:4 w/v ratio. The saps were rub-inoculated onto 2-leaf stage maize seedlings which were pre-dusted with 400-mesh carborundum powder (Sigma–Aldrich). As a parallel control, saps from mock-treated *N. benthamiana* leaves were rub-inoculated into maize seedlings. The newly emerged 4th leaves were collected at 10 dpi for investigating the silencing efficiency of *ZmTIRs* or scoring the SLB disease phenotype inoculated with *Cochliobolus heterostrophus*. All virus infection experiments were independently repeated at least three times.

### SLB Disease Scoring

The 4th leaves from CMV-silenced seedlings were evenly laid on two-layer filter papers in a clean black seedling tray. Holes were poked in the leaves with the inoculation needle. The spores of *Cochliobolus heterostrophus* grown on V8 agar medium were collected and diluted with rinsing solution to approximately 1 × 10^6^ spores per ml. 10 μL spore suspension was add to the holes of leaves. Disease symptom was evaluated after 2-4 days post inoculation (dpi) with calculating lesion areas of infected leaves using the software ImageJ.

## DATA AVAILABILITY STATEMENT

All relevant data can be found within the manuscript and its supporting materials.

## ACKNOWLEDGMENTS

We are grateful to Dr. Brian Staskawicz for sharing the *eds1* and *nrg1* mutants, Dr. Xiangxiu Liang for *adr1* and *nrg1/adr1* mutants, Dr. Johannes Stuttmann for *pad4* mutant, Dr. Li Wan for *epss* mutant, Dr. Yan Wang for *bak1* mutant, and Dr. Zhiyuan Yin for *sobir1* mutant. We are grateful to Dr. Keiko Yoshioka for providing the GCaMP3 transgenic *N. benthamiana* line, Dr. Yule Liu and Tao Zhou for providing CMV vectors for VIGS. We thank Haiyan Yu, Xiaomin Zhao and Sen Wang from SKLMT (State Key Laboratory of Microbial Technology, Shandong University) for the assistance in microimaging of LSCM analysis. This research is supported by grants from the National Natural Science Foundation of China (32072405 and 31871944), the National Key Research and Development Program of China (2022YFD1201802), the Key R&D Project of Shandong Province (2021LZGC022), the Strategic Priority Research Program of the Chinese Academy of Sciences (XDA0440000), and the Fundamental Research Fund of Shandong University.

## AUTHOR CONTRIBUTIONS

G.-F. Wang conceived and designed the research. Z-X. Kang performed most of the experiments and analyzed the data with help from Q.-D. Ge, C.-X. Tang, M.-L. Zhang, M. Yang, S. Liu, Y.-C. Dai, H. Zhang and K. Zang. G.-F. Wang wrote the manuscript. Z-X. Kang and S. Li edited the manuscript. All authors read and approved the final manuscript.

## SUPPORTING INFORMATION

**Figure S1. ZmTIR1- and ZmTIR2-mediated auto-HR is independent on BAK1 and SOBIR1.** GUS, CC_D21_ and ZmTIRs were transiently expressed in *N. benthamiana* mutants *bak1* and *sobir1*. A representative leaf (left) was shown at 3 dpi, and the same leaf (right) was cleared by ethanol. Ion leakage conductivity was measured at 60 hpi. Different letters (a-b) indicate significant differences (Average ± SE, n > 5, P < 0.05) between samples. Total proteins were extracted from agro-infiltrated *N. benthamiana* leaves at 36 hpi. The expression of different proteins was detected with anti-GFP. Ponceau S stained was used as an equal loading control for protein samples.

**Figure S2. The self-association of the ZmTIR mutants.** The NADase and BB-loop mutants of ZmTIR1 and ZmTIR2 still self-associate when detected by Co-IP assays. ZmTIR:EGFP and ZmTIR:Myc proteins were co-infiltrated into *N. benthamiana* and samples were collected at 40 hrs. Anti-GFP microbeads were used for immunoprecipitation (IP) and immunoblotting (IB) was performed by anti-GFP and anti-Myc antibodies.

**Figure S3. The prediction of IDRs in ZmTIRs and the subcellular localization of ZmTIR mutants.** (a) The IDR in ZmTIRs were predicted by PONDR. Boxed regions indicate the BB-loop region. (b) The subcellular localization of the mutants in affecting the putative NADase and BB-loop of ZmTIR1 and ZmTIR2. ZmTIRs and their mutant derivatives were infiltrated into *N. benthamiana* harboring the nuclear marker H2B:TaqRFP. Confocal images were taken at 36 hpi. The scale bar represents 50 µm.

**Table S1.** The primers used in this study.

