## Supplemental figures for "Maize TIR-only Proteins ZmTIR1 and ZmTIR2, but not ZmTIR3 Confer Auto-active Hypersensitive Response Likely by Forming Condensation"

### Slide 1
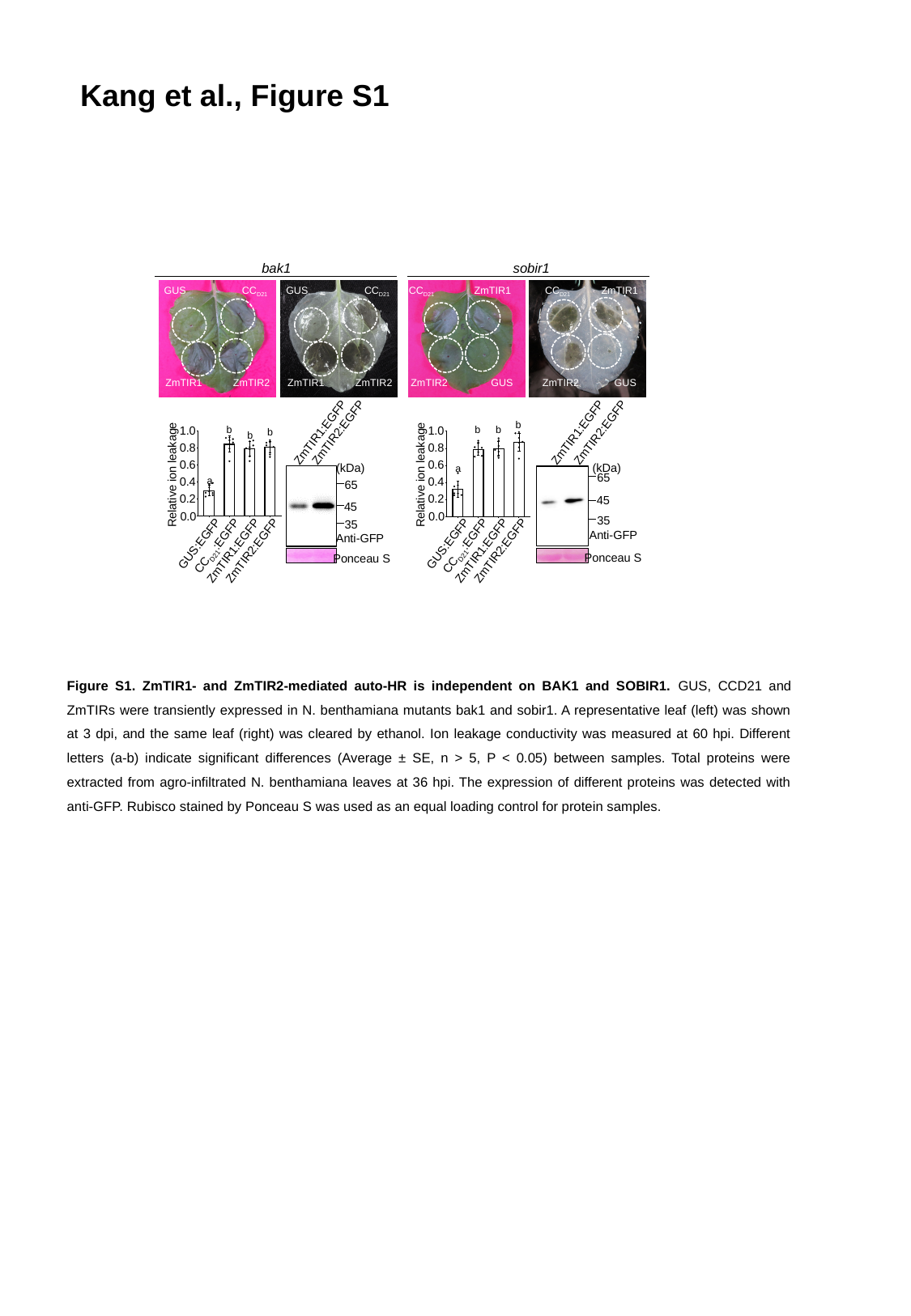

Kang et al., Figure S1
bak1
sobir1
GUS
CC_D21_
GUS
CC_D21_
CC_D21_
ZmTIR1
CC_D21_
ZmTIR1
ZmTIR1
ZmTIR2
ZmTIR1
ZmTIR2
ZmTIR2
GUS
ZmTIR2
GUS
1.0
b
b
b
0.8
0.6
Relative ion leakage
0.4
a
0.2
0.0
GUS:EGFP
CC_D21_:EGFP
ZmTIR1:EGFP
ZmTIR2:EGFP
b
b
1.0
b
0.8
0.6
Relative ion leakage
a
0.4
0.2
0.0
GUS:EGFP
CC_D21_:EGFP
ZmTIR1:EGFP
ZmTIR2:EGFP
ZmTIR1:EGFP
ZmTIR2:EGFP
ZmTIR1:EGFP
ZmTIR2:EGFP
(kDa)
(kDa)
65
65
45
45
35
35
Anti-GFP
Anti-GFP
Ponceau S
Ponceau S
Figure S1. ZmTIR1- and ZmTIR2-mediated auto-HR is independent on BAK1 and SOBIR1. GUS, CCD21 and ZmTIRs were transiently expressed in N. benthamiana mutants bak1 and sobir1. A representative leaf (left) was shown at 3 dpi, and the same leaf (right) was cleared by ethanol. Ion leakage conductivity was measured at 60 hpi. Different letters (a-b) indicate significant differences (Average ± SE, n > 5, P < 0.05) between samples. Total proteins were extracted from agro-infiltrated N. benthamiana leaves at 36 hpi. The expression of different proteins was detected with anti-GFP. Rubisco stained by Ponceau S was used as an equal loading control for protein samples.

### Slide 2
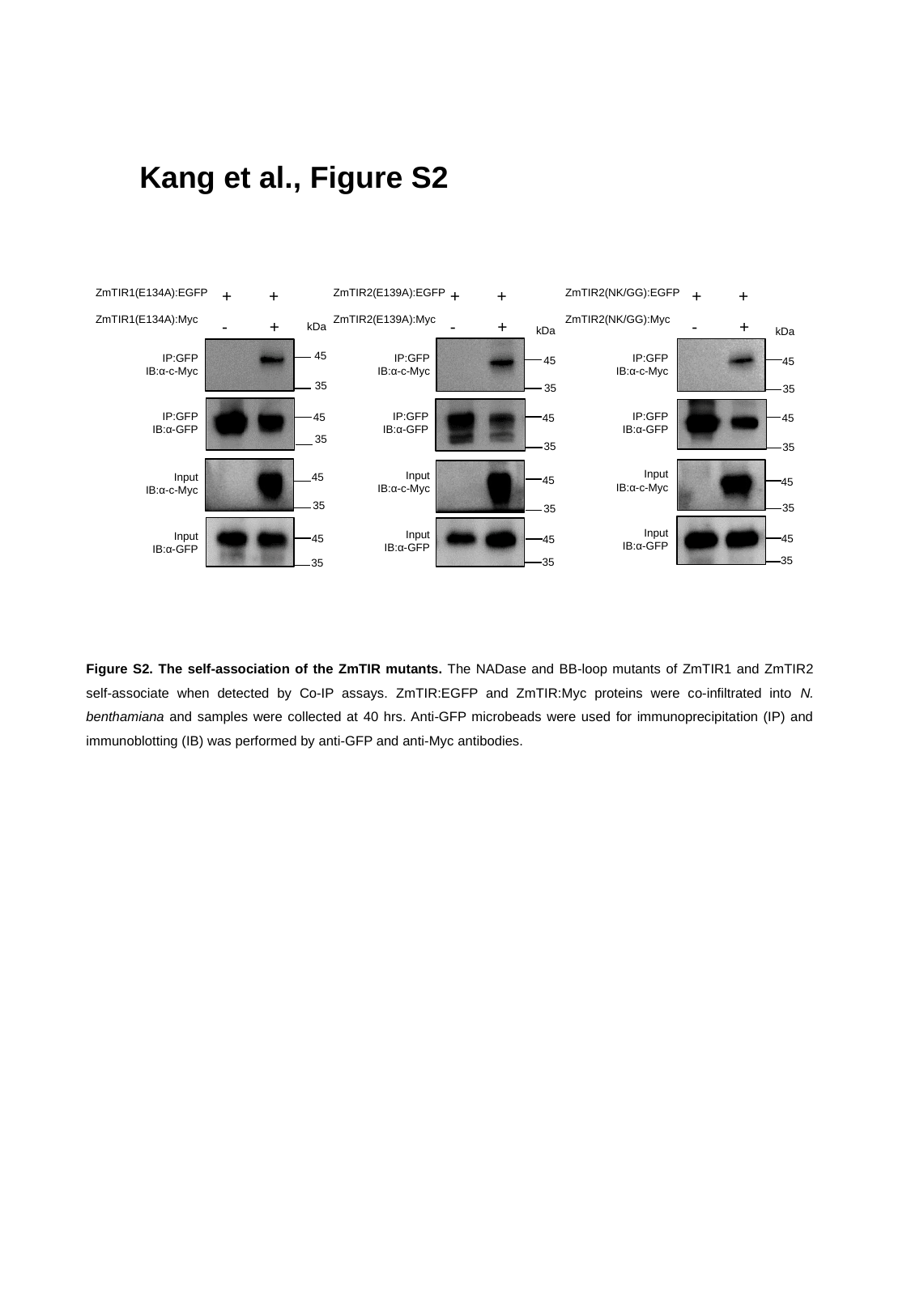

Kang et al., Figure S2
+ +
- +
+ +
- +
+ +
- +
ZmTIR1(E134A):EGFP
ZmTIR1(E134A):Myc
ZmTIR2(E139A):EGFP
ZmTIR2(E139A):Myc
ZmTIR2(NK/GG):EGFP
ZmTIR2(NK/GG):Myc
kDa
kDa
kDa
45
IP:GFP
IB:α-c-Myc
IP:GFP
IB:α-c-Myc
IP:GFP
IB:α-c-Myc
45
45
35
35
35
IP:GFP
IB:α-GFP
IP:GFP
IB:α-GFP
IP:GFP
IB:α-GFP
45
45
45
35
35
35
Input
IB:α-c-Myc
Input
IB:α-c-Myc
Input
IB:α-c-Myc
45
45
45
35
35
35
Input
IB:α-GFP
Input
IB:α-GFP
Input
IB:α-GFP
45
45
45
35
35
35
Figure S2. The self-association of the ZmTIR mutants. The NADase and BB-loop mutants of ZmTIR1 and ZmTIR2 self-associate when detected by Co-IP assays. ZmTIR:EGFP and ZmTIR:Myc proteins were co-infiltrated into N. benthamiana and samples were collected at 40 hrs. Anti-GFP microbeads were used for immunoprecipitation (IP) and immunoblotting (IB) was performed by anti-GFP and anti-Myc antibodies.

### Slide 3
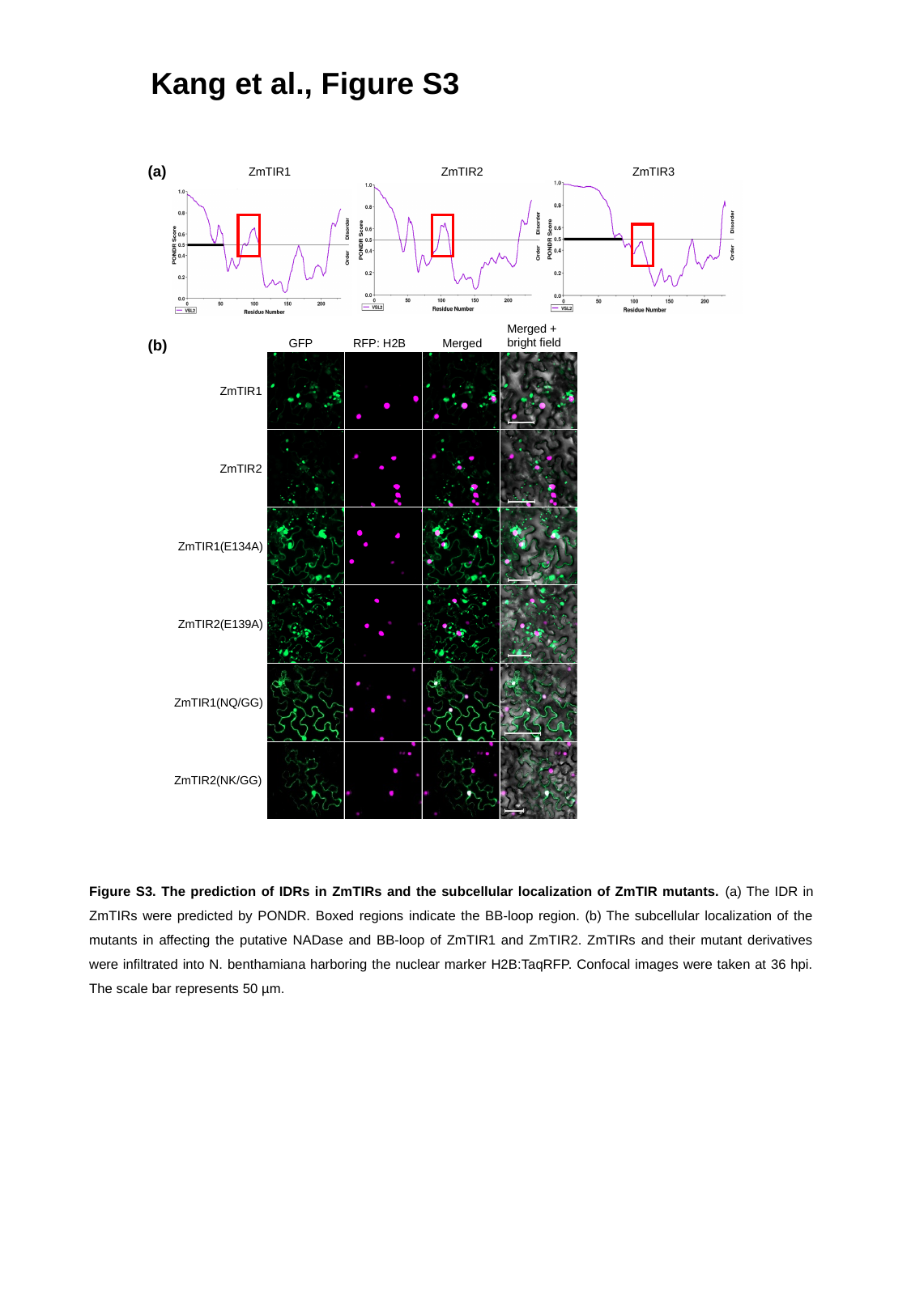

Kang et al., Figure S3
(a)
ZmTIR1
ZmTIR2
ZmTIR3
Merged +
bright field
(b)
GFP
RFP: H2B
Merged
ZmTIR1
ZmTIR2
ZmTIR1(E134A)
ZmTIR2(E139A)
ZmTIR1(NQ/GG)
ZmTIR2(NK/GG)
Figure S3. The prediction of IDRs in ZmTIRs and the subcellular localization of ZmTIR mutants. (a) The IDR in ZmTIRs were predicted by PONDR. Boxed regions indicate the BB-loop region. (b) The subcellular localization of the mutants in affecting the putative NADase and BB-loop of ZmTIR1 and ZmTIR2. ZmTIRs and their mutant derivatives were infiltrated into N. benthamiana harboring the nuclear marker H2B:TaqRFP. Confocal images were taken at 36 hpi. The scale bar represents 50 µm.
