## Supplemental table for "Maize TIR-only Proteins ZmTIR1 and ZmTIR2, but not ZmTIR3 Confer Auto-active Hypersensitive Response Likely by Forming Condensation"

Table S1. The primers used in this study.

| Primer name | Primer sequence（5'-3'） | Usage of the primers |
| --- | --- | --- |
| TIR1-F | ATGGCGTCGTCCGCGTTG | Amplication of *ZmTIR1* |
| TIR1-R | CAGCCTCGAAAGGATCATCTGGCG |  |
| TIR2-F | ATGGCGTCGTCCGCGCT | Amplication of *ZmTIR2* |
| TIR2-R | CAGCCTCGAAATGATCATCTGCTGGC |  |
| TIR3-F | ATGAGCAGCAGCGGGTCG | Amplication of *ZmTIR3* |
| TIR3-R | TCGCATGCTATCTATTTGTTCAATC |  |
| TIR1-qRT-F | TAACACCAGATCCGCCAAGT | qRT for *ZmTIR1* |
| TIR1-qRT-R | GATCAAACGCTTCACCCTCG |  |
| TIR2-qRT-F | TTAACACCAGGTCCGCCAAG | qRT for *ZmTIR2* |
| TIR2-qRT-R | GACTTGTACACGAGCCAGGA |  |
| TIR3-qRT-F | GCCTCGCCTACGACCCA | qRT for *ZmTIR3* |
| TIR3-qRT-R | TTCAATCCTCTCCATCACCGT |  |
| Zmactin-qRT-F | TACCATGTTCCCTGGGATTG | qRT for *Zmactin* |
| Zmactin-qRT-R | GTGGCGCAATCACTTTAACC |  |
| TIR1-E134A-MF | CTCCGCGCGCTCGCCT | Amplication of *ZmTIR1^E134A^* |
| TIR1-E134A-MR | AGGCGAGCGCGCGGAG |  |
| TIR2-E139A-MF | CTCCGCGCGCTCGCCT | Amplication of *ZmTIR2^E139A^* |
| TIR2-E139A-MR | AGGCGAGCGCGCGGAG |  |
| TIR1-C131A-MF | GACTTCGCCCTCCGCGAG | Amplication of *ZmTIR1^C131A^* |
| TIR1-C131A-MR | CTCGCGGAGGGCGAAGTC |  |
| TIR2-C136A-MF | CCGACTACGCCCTCCGCGA | Amplication of *ZmTIR2^C136A^* |
| TIR2-C136A-MR | TCGCGGAGGGCGTAGTCGG |  |
| T1/2-bl-MF | CGGGTCCGCTCCTTCCTGGACGGAGGAGGAGGAGGAAGCGGAGGAAGGCTCCA | Amplication of *ZmTIR1^bl^* and *ZmTIR2^bl^* |
| T1/2-bl-MR | TGGAGCCTTCCTCCGCTTCCTCCTCCTCCTCCGTCCAGGAAGGAGCGGACCCG |  |
| T1/2-NQNK/SV-MF | GCTCCTTCCTGGACAGCGTGTCCATGCGCCCCG | Amplication of *ZmTIR1^NQ/GG^* and *ZmTIR2^NK/GG^* |
| T1/2-NQNK/SV-MR | CGGGGCGCATGGACACGCTGTCCAGGAAGGAGC |  |
| TIR1/2-VIGS-F | GGGGTACCCGTTAACACCAGATCCGCCA | VIGS for *ZmTIR1* and *ZmTIR2* |
| TIR1/2-VIGS-R | CGGCTAGCTTCTACGGCATCAAGCCCTC |  |
| TIR3-VIGS-F | GGGGTACCCACCGTCTTGGCTGCCG | VIGS for *ZmTIR3*  VIGS for *ZmTIR3* |
| TIR3-VIGS-R | CGGCTAGCTGGTGCTGCCGCAGGC |  |
